# Long-term Connectivity between Spinal Cord Tissue Transplants and the Injured Phrenic Motor Network

**DOI:** 10.64898/2026.07.23.738908

**Authors:** Adam A. Hall, Lyandysha V. Zholudeva, Theresa Connors, Victoria Spruance, Tara Fortino, Kayla Schardien, Alessia Niceforo, Kimberly J. Dougherty, Liang Qiang, Michael A. Lane

## Abstract

Restoring vital motor functions after spinal cord injury (SCI) remains a central challenge in neuroscience and regenerative medicine. Cervical SCI can cause life-threatening respiratory deficits by damaging the phrenic motor network that controls the diaphragm. Cellular transplantation offers a viable means to improve function by providing new neurons that can relay supraspinal drive to denervated spinal phrenic networks, yet the long-term stability of transplants is poorly defined. Here, we examine donor-host neuronal synaptic connectivity in a respiratory model of cervical SCI, 1-year post-transplantation in adult rats. Embryonically-derived spinal cord tissue was transplanted into the lesion cavity one-week post-SCI, and transplant integration and diaphragm function were assessed at 1-month and 1-year post-transplantation. At 1-month, transplant-recipients exhibited significantly greater diaphragm output than injured, vehicle control animals. The extent of recovery at 1-year, however, was significantly less, coinciding with anatomical changes in graft neuronal density and donor-host connectivity, revealed by transneuronal tracing (pseudorabies virus). These results demonstrate that embryonic spinal cord transplants can improve phrenic motor activity after cervical SCI, but that long-term efficacy may be limited by reduced donor-host connectivity.

**Significance Statement:** Cell transplantation can repair injured spinal cord circuits, but whether donor-host connections persist long term remains unclear. Using a rat model of cervical spinal cord injury, we show that embryonic spinal cord transplants improve diaphragm activity and integrate with the injured phrenic motor network at early time points, but these benefits decline by 1 year after transplantation. This loss of functional recovery is accompanied by reduced transneuronal labeling of donor neurons and changes in graft tissue composition. These results provide important proof of principle that transplant-host connectivity can be evaluated over extended survival times and identify long-term stability of donor-host integration as a critical challenge for achieving durable respiratory repair after spinal cord injury.

## Introduction

Spinal cord injury (SCI) devastates the lives of tens of thousands of individuals in the US each year, often carrying with it fatal consequences (NSCISC, 2025). The majority of injuries occur at cervical levels resulting in both denervation and direct damage to networks at that level. Among those is the phrenic motor network, which controls the diaphragm, a primary respiratory muscle (Fogarty and Sieck, 2020). While some spontaneous plasticity typically occurs sub-acutely after cervical injury, it is limited, and people remain at risk of life-threatening complications such as respiratory insufficiency and pneumonia (DeVivo et al., 1989; Kirshblum et al., 2021; Locke et al., 2022). Although there is growing effort to therapeutically enhance plasticity and lasting recovery (e.g., neural stimulation or activity-based therapies), these approaches do not treat the underlying damaged anatomy. In contrast, cell therapies have been identified as translationally relevant strategies for spinal cord repair (Reier et al., 1986; Ogawa et al., 2002; Lu et al., 2012). Cell therapies limit secondary damage through attenuated inflammation and host tissue loss while restoring tissue integrity and providing a bridge for host axonal growth along with cell migration. These treatments also provide a source of donor neurons and glia to build new neural networks that contribute to repair and recovery (Reier, 1985; Zholudeva and Lane, 2019b; Hall et al., 2022). Previous studies by our team (White et al., 2010; Lee et al., 2014; Spruance et al., 2018; Zholudeva et al., 2018; Zholudeva et al., 2024) and others (Li et al., 2015a; Li et al., 2015b; Lin et al., 2017) have shown that transplanting neural progenitor cells (NPCs) can repair the damaged phrenic network and improve breathing after cervical SCI. While a number of different donor cell sources have now been tested, transplantation of embryonic spinal cord tissue provides an excellent ‘proof-of-concept’ for studying spinal cord repair. These transplants contain the full complement of cellular and non-cellular elements necessary to reconstruct neural architecture and promote tissue repair. Among these are pro-reparative glia (Smith and Silver, 1988; Jin et al., 2011; Goulao et al., 2019) and spinal interneurons (Zholudeva et al., 2017; Zholudeva et al., 2021b; Kathe et al., 2022; Zholudeva and Lane, 2022a) that are known to contribute to circuit reconstruction, plasticity and recovery following injury. As a deeper understanding of the reparative capacity of these cells emerges, transplanting their neural progenitors has become a focus of cell therapies for spinal repair.

Neuronal progenitors transplanted into the injured spinal cord have been shown to be innervated by the host and extend donor axons into the injured host, synaptically integrating to form novel networks that can improve motor and sensory function (Bonner et al., 2011; Lu et al., 2012; Adler et al., 2017; Dulin et al., 2018; Kumamaru et al., 2018; Kumamaru et al., 2019; Ceto et al., 2020; Sanchez-Martin et al., 2025). Most prior studies, however, have focused on early integration within weeks to months after transplantation, leaving it unclear whether newly established donor-host synapses persist, or are lost (e.g., via remodeling, pruning or degeneration) over extended survival periods. This represents a critical gap in our understanding of transplant-mediated circuit repair. The present study addresses this gap by examining long-term donor-host connectivity of embryonic spinal cord transplants with the injured phrenic motor network following cervical SCI.

Using transsynaptic neuroanatomical tracing (Fortino et al., 2022) and functional assays (Lane et al., 2012) at 1-month and 1-year time points, we aimed to determine whether early graft-mediated integration and diaphragm recovery are sustained or lost a year following a lateralized cervical (C3/4) level contusion injury in adult rats. Consistent with previous work, transplanted tissue survived, and donor neurons became integrated with the host phrenic network ipsilateral to injury, but the extent of connectivity was reduced over time. In parallel, there was a decline in recovered diaphragm activity. The present work demonstrates for the first time that there is a progressive loss of integration over long periods of time that correlates with functional decline.

## Materials and Methods

Animal experiments were performed in compliance with the NIH Guidelines for Animal Care and Use of Laboratory Animals and approved by Drexel University College of Medicine Institutional Animal Care and Use Committee. Adult, female Sprague-Dawley (SD) rats (N=69; 225-300g; Envigo) were divided into experimental (transplant recipients) and control groups (naïve/uninjured, injured only, and injured, vehicle treated). These four groups were studied for 1 month (1-M) and 1 year (1-Y) post-injury and transplantation. Naive animals were age-matched to injured animals; approximately 4 months (N=10) and >13 months (N=15). All other animals were acclimated to the facility for at least 1 week after arrival before undergoing C3/4 lateralized contusions. Injury controls were divided into injury only and injured, vehicle-treated animals. Injury only animals received no interventions after the initial injury (N=4 at 1-M; N=10 at 1-Y). Vehicle treated animals underwent intraspinal injections with Hanks’ balanced salt solution (HBSS; N=6 at 1-M; N=4 at 1-Y). Transplant recipients received intraspinal (intralesional) injections of dissociated fetal spinal cord (FSC) tissue derived from transgenic GFP-expressing SD time-mated dams (N=9 at 1-M; N=10 at 1-Y). Three days prior to terminal electrophysiology pseudorabies virus (PRV) was topically applied onto the ipsilateral-to-injury hemidiaphragm muscle (N=69). Following terminal electrophysiology all tissues were perfusion-fixed. A schematic diagram of the methods is depicted in Figure 1.

**Figure 1.**
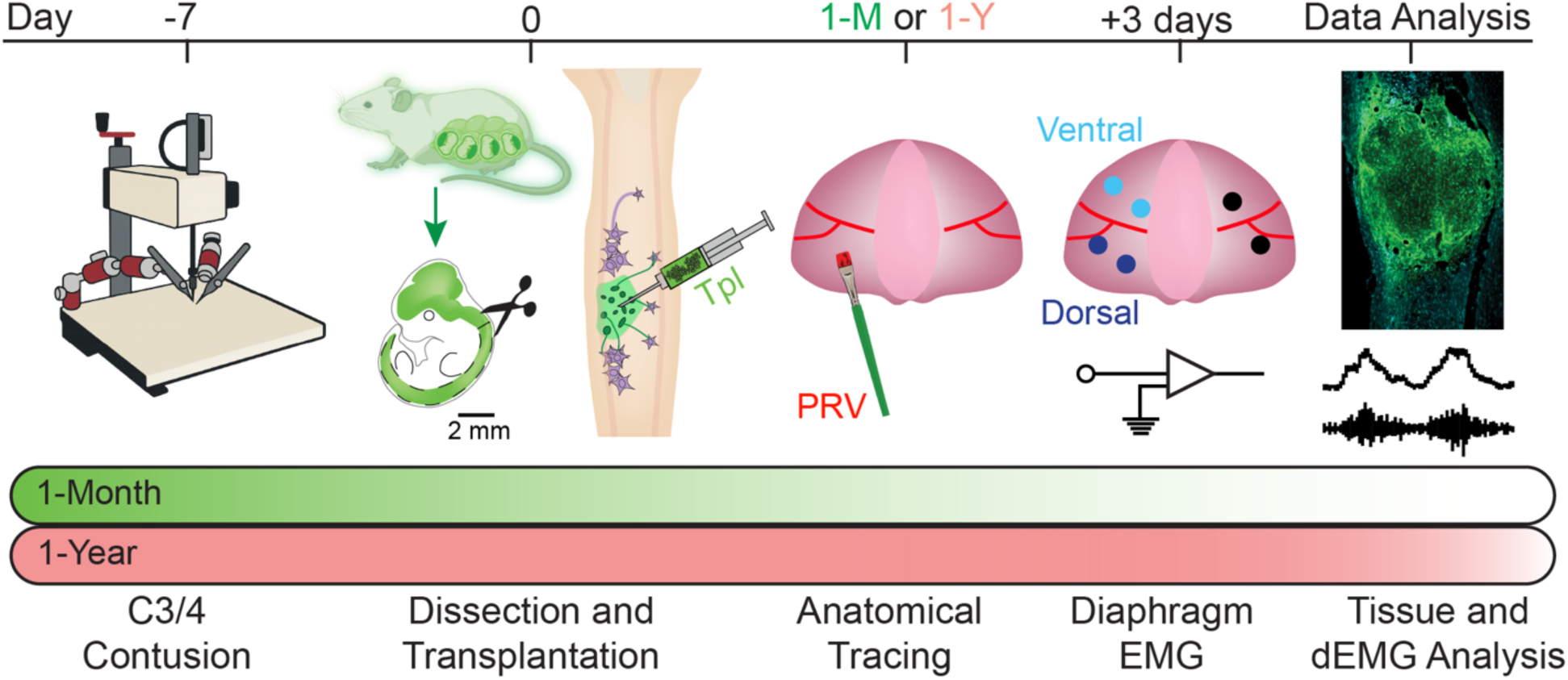
Schematic of the experimental design and timeline. Fourteen days prior to transplantation, transgenic eGFP-expressing Sprague-Dawley rats were time-bred to generate GFP-positive embryos for embryonic day 13.5 spinal cord dissections. Seven days prior to transplantation (-7 days), animals received a lateralized C3/4 contusion injury using an IH impactor. At the time of transplantation (day 0), embryos were collected, spinal cords were dissected, and mechanically dissociated as previously described (Zholudeva et al., 2021a). Cells were delivered into the lesion cavity using a stereotaxic frame and a Hamilton syringe with custom beveled, 30-gauge needle. For anatomical tracing, Pseudorabies virus (PRV), a transsynaptic retrograde tracer, was applied topically to the ipsilateral-to-injury hemidiaphragm (1-M or 1-Y), 3 days prior to terminal experiments. Diaphragm electromyography (EMG) recordings were obtained at terminal timepoints (+3 days) by inserting electrodes into dorsal and ventral regions of the ipsilateral hemidiaphragm and into the contralateral hemidiaphragm. Diaphragm EMG analysis, tissue processing and immunohistochemistry (Data Analysis) were used to quantify anatomical and functional outcomes across experimental groups.

### Spinal cord injury

Lateralized contusions at the C3/4 level have been described previously (Lane et al., 2012; Spruance et al., 2018; Zholudeva et al., 2018). Briefly, animals were anesthetized with sequential injections of xylazine (10 mg/kg, subcutaneous (s.q.); Anased®; Akron Animal Health, Lake Forest, IL) and ketamine (120 mg/kg, intraperitoneal (i.p.); KetaVed®; Vedco, St. Joseph, MO), 10 minutes apart, to achieve a surgical plane of anesthesia. Animals were laid prone, skin overlying cervical musculature prepared for surgery, and an incision made through the skin and musculature over the 2^nd^ to 5^th^ cervical vertebrae. The muscles were retracted and a partial laminectomy of C3 and rostral C4 was performed to expose the dorsal spinal cord. A left-lateralized contusion injury was performed at C3/4 (between C3 and C4 rootlets) using the Infinite Horizon pneumatic impactor (Precision Systems & Instrumentation, Fairfax Station, VA; Figure 1), with a preset impact force of 200 kilodynes (KD; zero dwell time). The extent of subdural bleeding, spinal bruising, and respiratory consequences (e.g. arrest) were noted. Animals experiencing respiratory arrest were immediately intubated then placed on a rodent mechanical ventilator. All ventilated animals were given a respiratory stimulant, doxapram hydrochloride (Dopram®, 5mg/kg, s.q.; West-Ward, Eatontown, NJ) to assist with weaning from ventilation. Animals were removed from ventilator support within an hour following injury, and those animals that could not regain voluntary control of breathing by this time were excluded from the study. Muscles were sutured (4-0 or 5-0 Vicryl) and skin closed with wound clips. Animals were given atipamezole hydrochloride (Antisedan®; 2.5 mg/kg, s.q.; Astatech, F20658) to counteract anesthesia, and Lactated Ringers (5ml, s.q.) for hydration. Animals were group housed and provided with care 3-times daily following the injury until animals were able to eat and drink from suspended sources in the cage. Care included cleaning, bladder expression, feeding with a calorie-dense Nutrical solution (Vetaquinol, EVS2747) and lactated ringer injections until they were able to reach a suspended water bottle.

### Spinal cord tissue transplantation

Experimental protocols for isolating the developing spinal cord tissue have been described previously (Zholudeva et al., 2021a). Embryos (embryonic day ∼13.5; E13.5) were extracted from time-mated SD-eGFP rats. The embryonic spinal cords were dissected in HBSS in a laminar flow hood. Dissection of spinal cord tissue ensured the exclusion of meninges, dorsal root ganglia, and any brain or brainstem tissue (Bonner et al., 2013; Zholudeva et al., 2021a). Although this donor tissue is often referred to as “fetal spinal cord”, this is a misnomer, as the tissue is derived from an embryonic developmental stage (Altman and Katz, 1962). Once dissected, pieces of spinal cord tissue were mechanically dissociated into a suspension in sterile HBSS. Donor tissue suspensions primarily comprise small tissue pieces and cell clusters, rather than single cells. Cell number and viability was determined before transplantation (Cellometer, Auto T4, Nexcelom Biosciences; trypan blue dilution), and viability was greater than 75% in all recipients.

One-week post-contusion, animals were re-anesthetized, and the laminectomy and injury site were re-exposed. Scar tissue overlying dura was gently removed, and a small opening was first made through the dura and dorsal spinal cord with an 18-gauge needle to then allow the beveled injection needle to access the lesion cavity. The transplant tissue (1.0×10^6^ cells in 10µL of HBSS) or vehicle (10µL of HBSS) was stereotaxically injected using a gastight Hamilton syringe with a custom 30-gauge needle (Leur-lock, 45-degree bevel) over approximately 2 minutes. Needle was kept in place for 30 seconds prior to slow withdrawal. Muscles were sutured and skin was closed with wound clips. The animals received post-op medication and returned to care (as described above). Given that the donor and host tissue were both from the SD strain of rats, immunosuppression was not required.

### Neuroanatomical Tracing

Transneuronal tracing with PRV used in these experiments has been described in detail previously (Zholudeva et al., 2018; Fortino et al., 2022). Briefly, 72-75 hours prior to terminal experiments (1-M and 1-Y time-points) animals were anesthetized with isoflurane (induced with 4% in 100% oxygen, maintained 2-2.5%, flow rate 2L/min; South Medic In., Barrier, Ontario, Canada). Animals were prepared for aseptic abdominal surgery and received a laparotomy to expose the diaphragm. The retrograde, transsynaptic tracer (PRV614 Bartha strain; promotes expression of red fluorescent protein; 50 µL, titer 2.2×10^9^ PFU/mL; University of Florida) was topically applied to the ipsilateral-to-injury hemi-diaphragm using a fine-hair paint brush, as previously described (Lane et al., 2008). After application, the abdominal wall was sutured, the skin was closed with wound clips, and the animals received post-op medication and care.

### Terminal Diaphragm Electromyography

At the end of the study, animals were surgically anesthetized for terminal electrophysiology with xylazine/ketamine as described above (n=69). Fur over the neck was shaved, animals placed in a supine position on a heating pad and had a Mouse-Ox Plus® neck clip placed to measure oxygen saturation (SpO_2_) and heart rate for the duration of the recording (STARR Life Sciences Corp., Oakmont, PA, 015029). Once SpO_2_ was stable >93%, the diaphragm was re-exposed, and tungsten electrodes (PFA-coated tungsten wire; A-M Systems, Inc., Sequim, WA) were placed in the diaphragm for bipolar electromyography (EMG): two electrodes were implanted in the left ventro-medial costal diaphragm (ipsilateral to injury/transplant), two in the left dorsal costal diaphragm, and two in the right medial diaphragm (contralateral to injury/transplant; see Figure 1 Diaphragm EMG). All recordings were amplified (A-M Systems Differential AC amplifier, Model 1700), digitized (Power3 1401 digitizer; Cambridge Electronic Design, Ltd. [CED], Cambridge, UK), and recorded using Spike2 software (version7; CED). Diaphragm traces were rectified (DC bias removed with 0.1-sec window width) and integrated (T=0.03 sec, time constant).

All recordings are made during spontaneous breathing. Once electrodes were placed, the animals’ airways were briefly sealed (10 seconds; inspiratory restriction, IR) to elicit strong inspiratory drive and diaphragm contraction. This was followed by 10 minutes baseline eupneic breathing (room air), 5 minutes of hypercapnic challenge via a nose cone (inhaled air with 7% CO_2_ balanced in N_2_; ∼2.5L/min), 5 minutes rest (room air), then 5 minutes of hypoxic challenge (10% O_2_ balanced in N_2_; ∼2.5L/min). After another 5-minute recovery period (room air) the animals were given 3 x 30 second inspiratory restriction (IR) challenges (nose and mouth sealed to prevent spontaneous inspiration) with 2 minutes recovery in between, to assess increased diaphragm activity. Animals were then euthanized and tissues perfuse-fixed.

Average diaphragm EMG amplitudes were calculated from integrated signals during baseline and each challenge (defined as the last 2 minutes of hypercapnia or hypoxia). For inspiratory restrictions, the breath exhibiting the largest diaphragm amplitude was used for analysis. The response to each challenge was calculated by dividing the diaphragm amplitude during the challenge (hypercapnia, hypoxia, or inspiratory restriction) by the baseline and presented as a ratio.

Breathing frequency was quantified from the same recordings by counting individual breaths over a 40 second sampling window and comparing values across baseline, hypercapnic, and hypoxic challenges. Data were analyzed across three groups: naïve (uninjured) animals, injured animals (injury only or injury with vehicle treatment), and injured animals that received transplants at 1-M and 1-Y timepoints. There were no significant differences between animals that were injured only and vehicle treated (1-Month hypercapnia p=0.84, hypoxia p=0.525, inspiratory restriction p=0.436; 1-Year hypercapnia p=0.602, hypoxia p=0.918, inspiratory restriction p=0.087), thus data were combined for statistical analyses and plotted together, with individual animals color-coded to indicate group identity.

### Histology

Animals were intracardially perfused with 0.9% saline (250ml) and 4% paraformaldehyde (in 0.1M phosphate buffered saline; 500ml). The dissected spinal cords were post-fixed in 4% for >12 hours. The cervical cord was cryo-preserved in sucrose solutions, and frozen in M1 embedding matrix. Frozen tissues were sectioned on slide (20 µm thickness, longitudinal) using a Leica cryostat. Immunohistochemistry was conducted using primary antibodies against PRV (1:10,000; courtesy of Dr. Lynn Enquist at Princeton University or 1:500; Encor Biotechnology Inc), Serotonin (1:500; 20079 Immunostar, Hudson, WI), Neuronal nuclei protein (NeuN; MAB377, 1:500; EMD Millipore), Iba1 (RPCA-IBA4,1:500; Encor Biotechnology Inc.). Secondary antibodies used were: Donkey-anti-Mouse 647 (1:200, Invitrogen, A32787), Donkey-anti-Chicken, 488 (1:200, Jackson Immunoresearch, 703-545-155), Donkey-anti-Goat, 594 (1:200, Jackson Immunoresearch, 705-585-003) 647 (1:200, Jackson Immunoresearch, 705-605-003), Donkey-anti-Rabbit, 594 (1:200, Jackson Immunoresearch, 711-585-152). All images were taken using Zeiss Imager M2 with Apotome.2 and Zen Blue software (Carl Zeiss, Oberkocken, Germany), or with a Leica DM6 B Thunder Imager and LasX Software (Leica, Wetzlar, Germany).

### Anatomical Quantification

All anatomical analyses were performed on images obtained from a Leica (Thunder Imaging System; LIF format). Quantification of PRV-positive cells was performed manually, counting all PRV-positive neurons in every 6^th^ section (120 microns apart). The number of PRV-positive donor neurons is represented as a proportion of labeled phrenic motor neurons (primary labeling). The density of NeuN, 5-HT, and Iba1 immunolabeling within the transplant (GFP+) area was measured using ImageJ, with GFP labeling used to define the region of interest. For serotonergic axons, 5-HT immunoreactivity was quantified as mean pixel intensity within defined regions of interest under identical imaging conditions, consistent with prior approaches assessing ventral horn serotonergic innervation (Starkey et al., 2012; Bartus et al., 2014).

Quantitative measurements of transplant and lesion volume were made in transplant recipients using ImageJ. GFP+ immunolabeled area, and injury site area (comprising both GFP+ transplant and any regions absent of tissue), were measured in ImageJ. The proportion of injury site filled with transplanted tissue is represented as a percentage. Serial area measurements (every 6^th^ section) were used to obtain an estimated transplant area for each animal, as described previously (Lu et al., 2017).

### Statistical Analyses

All statistical analyses were performed in Origin 2023b. Shapiro-Wilk testing determined normality and whether parametric or non-parametric hypothesis tests were used. All anatomical comparisons utilized parametric two sample t-tests or non-parametric Mann-Whittney Tests. In tests where 3 or more conditions were tested, parametric one-way ANOVAs or non-parametric Kruskal-Wallis ANOVAs were used with Tukey, Dunn’s, 2 sample-T, and Mann Whitney tests used as post-hoc tests. During an analysis where one factor is a between-groups design while the other is a within-groups design, Two-Way Mixed Measures ANOVAs were used with one-way ANOVAs or Kruskal-Wallis ANOVAs used as post-hoc tests. Pearson or Spearman correlations (parametric and non-parametric, respectively, depending on data distribution) were used to assess relationship between datasets. Data sets were subjected to an outlier Grubb’s test if the n-number exceeded 10, or a Q-test if the n-number was 10 or fewer. To ensure a strict standard for data removal, the significance threshold for outlier tests was set at 0.01. All hypotheses tested used a p-value of 0.05 as a significance threshold.

## Results

### Transplant Mediated Diaphragm Recovery is Reduced Over Time

Given the lateralized damage to the phrenic network on the left side, the primary focus of our analyses was on the ipsilateral hemi-diaphragm. The dorsal region is also more affected by this SCI as it is innervated by the more caudal phrenic motor neuron pool (caudal to injury and denervated). In contrast, the ventral region of the ipsilateral diaphragm may be less affected as it is innervated by the more rostral portion of phrenic motor pool (C3, Figure 1) (Laskowski and Sanes, 1987). Given there were no significant differences between injured animals, whether or not they received vehicle treatment, the results from these animals were combined and called ‘injured’ for all statistical analysis and graphs shown.

Baseline diaphragm activity (spontaneously breathing room air) in 1-M transplant recipients was significantly greater than age-matched naïve animals in the ipsilateral dorsal (p=0.012) and ventral (p=0.046) diaphragm (Supplementary Figure 1). In contrast, mean baseline ipsilateral diaphragm activity was reduced in 1-Y transplanted compared to naïve animals. While there was no difference in ipsilateral ventral muscle activity between groups, ipsilateral dorsal diaphragm activity in 1-Y transplant recipients was significantly less than age-matched naïve animals (p=0.002). Contralateral diaphragm activity in both injured animals and transplant recipients at 1-M was greater than time matched naïve animals (p=0.032 and 0.003, respectively; Supplementary Figure 1D). By 1-Y, however, there was no difference in contralateral diaphragm activity between groups. Baseline diaphragm activity in 1-M transplant recipients was greater that 1-Y recipients (p=0.003; Supplementary Figure 1B).

Animals were exposed to 3 respiratory challenges - hypercapnia (7% CO2), hypoxia (10% O2), and inspiratory restriction – to increase respiratory drive and evaluate diaphragm activity (Road et al., 1986; Shiota et al., 2004; Rana et al., 2017; Zholudeva et al., 2018). The response to hypercapnia (p=0.035) and inspiratory restriction (p=0.003) in injured animals at 1-M was significantly less than in age-matched naïve controls (Figure 2). However, there was no difference between naïve and injured animals in their response to hypoxia at 1-M (p=0.477; Figure 2C). Transplant recipients at 1-M displayed significantly greater responses to hypoxia (dorsal hemidiaphragm p=0.028; Figure 2C) and inspiratory restriction (dorsal hemidiaphragm p=0.011; Figure 2D and ventral hemidiaphragm p=0.037; Supplementary Figure 2) challenges than injured animals. These data show that transplant recipients have an improved response to respiratory challenge compared to injured animals at this early time point, which is comparable to age-matched uninjured controls.

**Figure 2:**
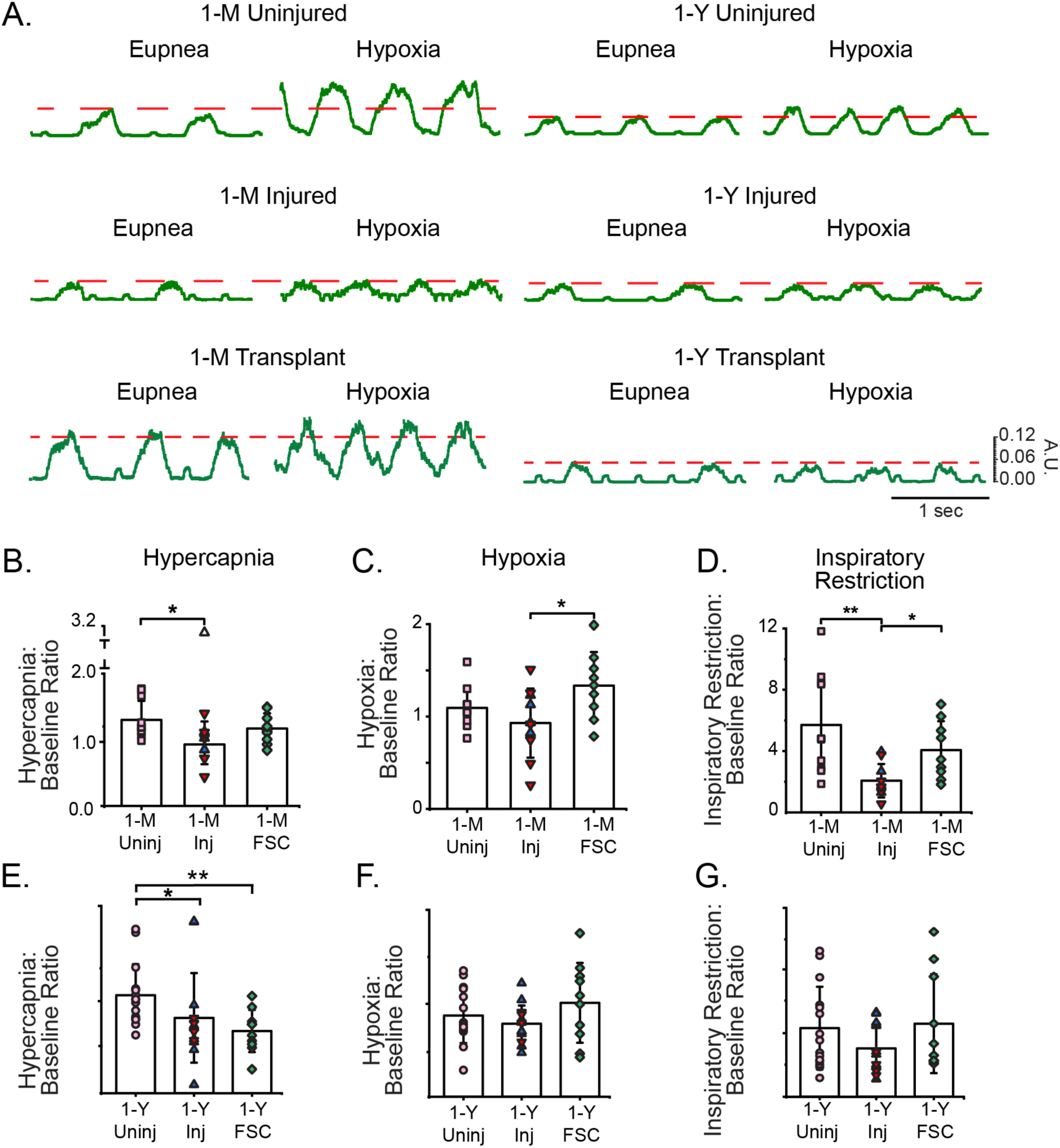
Ipsilateral dorsal diaphragm responses to challenge differ between 1-M and 1-Y cohorts. (A) Example dEMG traces illustrating eupneic (baseline) activity and the response to hypoxic challenge in 1-month (1-M) and 1-year (1-Y) cohorts. (B-D) Challenge-evoked dEMG amplitudes for the 1M cohort (naïve: N=10; injury only: blue, N=4; vehicle-treated: red, N=6; transplant-recipients: FSC, N=9), normalized to baseline. (B) Hypercapnia: naïve animals show a significantly greater response than injured cohort (p=0.0351), (C) Hypoxia: transplant recipients showed significantly enhanced response compared to injured controls (p=0.028) (D) Inspiratory restriction: both naïve animals (p=0.0031) and transplant recipients (p=0.0114) responded significantly greater than injury controls. (E-G) Challenge evoked dEMG amplitudes for the 1Y cohort (naïve: N=15; injury only: blue, N=10; vehicle, red N=4; and transplant-recipient: FSC, N=10). (E) Hypercapnia: both 1Y transplanted (p = 0.0083) and injured (p = 0.0276) animals show a significantly weakened response to hypercapnic challenge compared to 1-Y naïve animals. No other statistically significance differences were found in either response to hypoxia (F) or inspiratory restriction (G). * p<0.05; ** p<0.01.

In contrast to 1-M, transplant recipients at 1-Y showed an attenuated response to challenges, which more closely resembles that seen in injured animals (Figure 2E-G). 1-Y transplant recipients (p=0.008) and injured animals (p=0.028) had significantly lower responses to hypercapnic challenge than naïve animals (Figure 2E). There were no differences in response to hypoxia and inspiratory restriction challenges between naïve, injured, and 1-Y transplant recipients.

Breathing frequency calculated from EMG recordings also offers valuable insights into the consequences of injury and transplantation from 1-M to 1-Y. Although change in frequency across conditions was mostly comparable, transplant recipients at 1-M showed greater recovery in the ability to respond to hypercapnic challenge by breathing faster compared with time-matched vehicle treated animals (p=0.007) (Supplementary Figure 3). However, this response to challenge was lost in animals at 1-Y (p=0.506) (Supplementary Figure 3), providing further evidence for the loss of recovery 1 year post transplantation. Consistent with clinical observations, where people living with SCI exhibit an impaired response to respiratory challenge and can undergo respiratory arrest (Berlowitz et al., 2016), this was seen in a subset of injured animals at 1-M and 1-Y, but only in transplant recipients at 1-Y.

### Transplant Survival and Neuronal Density

The extent of transplant survival and lesion cavitation was assessed at 1 month (1-M) and 1 year (1-Y) post-transplantation (Figure 3A, B). Total donor transplant area did not differ significantly between 1-M and 1-Y recipients, despite increased inter-animal variability at the 1-Y time point (Figure 3C). In contrast, the proportion of the lesion cavity occupied by donor tissue was significantly greater at 1-Y compared with 1-M (p=0.003; Figure 3D), indicating progressive filling of the lesion site over time. These measures were used to examine relationships between transplant size, additional anatomical features (e.g., donor neuronal integration), and functional outcomes. Correlational analyses revealed no significant relationship between transplant size and diaphragm electromyographic activity at either time point (Supplementary Figure 4).

**Figure 3.**
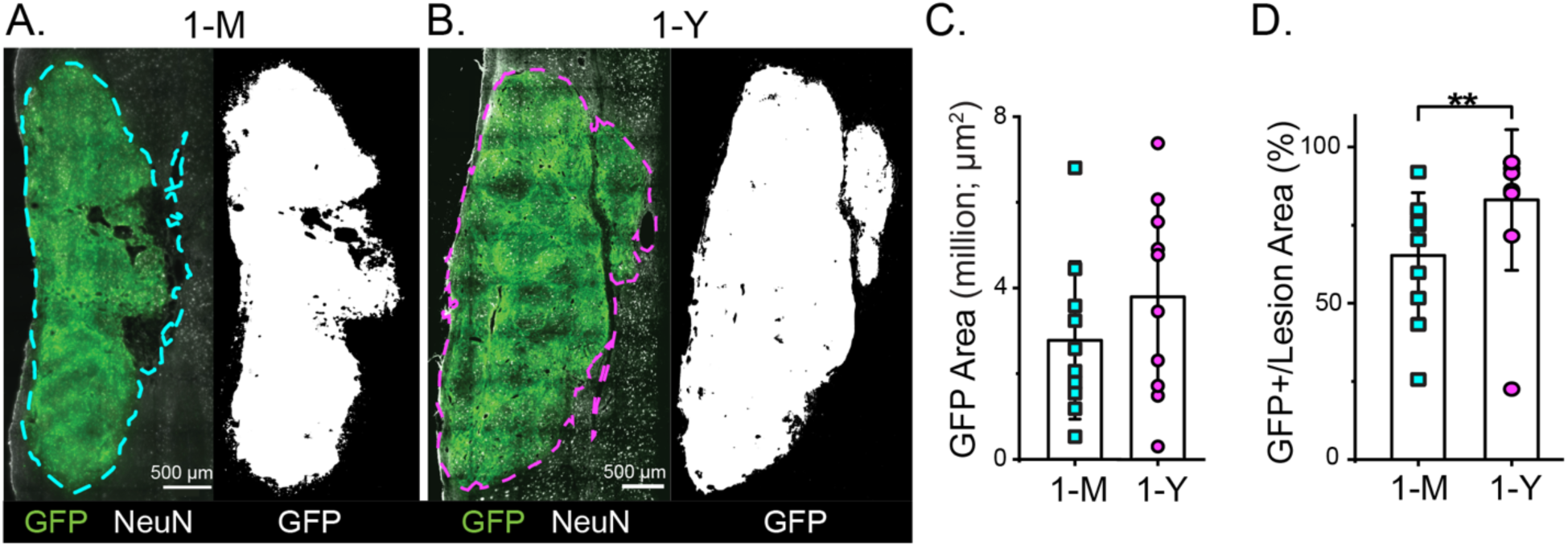
**Transplant growth and lesion cavity filling increase over extended survival times**. (A-B) Representative images of GFP+ donor cell grafts within the lesion cavity at 1 month (1-M, A) and 1 year (1-Y, B) post-transplantation. Dashed outlines indicate the lesion outline (left) and threshold set to GFP signal used for quantification (right). (C) Quantification of total GFP+ transplant area shows heterogeneity across animals, with some 1-Y grafts exhibiting larger areas; however, the overall difference between 1-M and 1-Y groups was not significant (2-sample t-test, p=0.289). (D) Percent cavity fill (GFP+ area normalized to total lesion area) was significantly greater in 1-Y animals compared with 1-M transplant recipients (Mann-Whitney Test, p=0.0029), indicating progressive graft expansion within the lesion cavity. * p<0.05; ** p<0.01.

To assess changes in neuronal content within transplanted tissue over time, neuronal density was quantified as NeuN+ signal normalized to total transplant area at 1-M and 1-Y post-transplantation (Figure 4A, B). Neuronal density within the transplant was significantly greater at 1-M compared to 1-Y (p=0.001; Figure 4C), indicating a reduction in NeuN+ cellular content over extended survival. Despite this difference, neuronal density at 1-M was not significantly associated with diaphragm EMG responses during hypercapnic, hypoxic, or inspiratory restriction challenges (Figure 4D). In contrast, at 1-Y, neuronal density showed a positive relationship with diaphragm EMG responses across respiratory challenges, reaching a strong trend during hypercapnia (r=0.804, p=0.054; Figure 4E).

**Figure 4.**
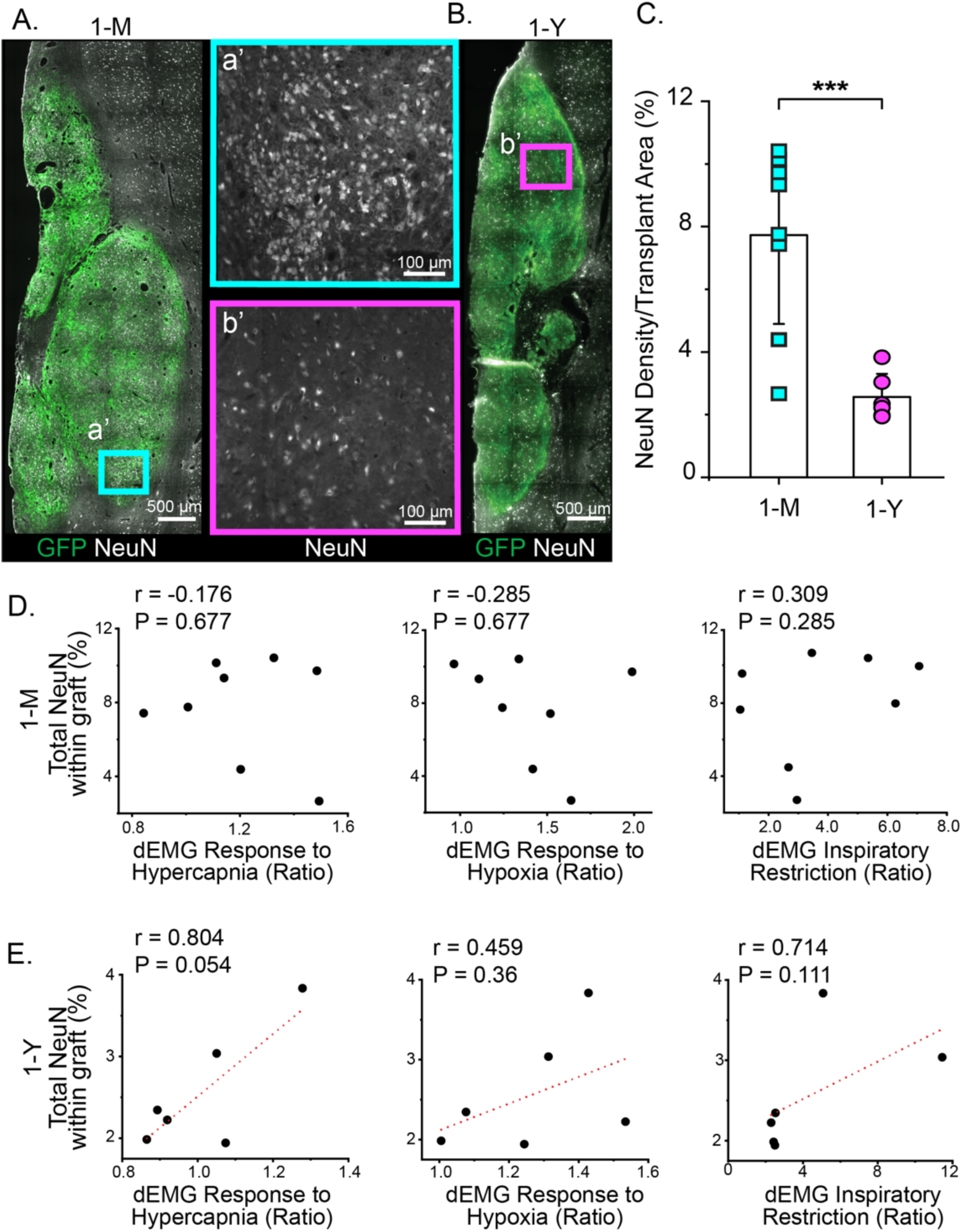
Transplant NeuN labeling decreases over time and its relationship with diaphragm EMG (dEMG) output differs between early and late survival times. (A-B) Representative images of NeuN+ neurons within GFP+ grafts at 1-month (1-M) and 1-year (1-Y) post-transplantation. Insets (a’, b’) show higher-magnification views of neuronal distribution within the graft. (C) NeuN+ labeling normalized to transplant (GFP+ labeling) area, was significantly lower in 1-Y grafts compared with 1-M grafts (2-sample t-test, p=0.001). (D) In 1-M animals, NeuN-labeling did not correlate with dEMG response during hypercapnia (Pearson Correlation, r=-0.176, p=0.677), hypoxia (r=-0.285, p=0.677) or inspiratory restriction (r=0.309, p=0.285). (E) In 1-Y animals, NeuN-labeling was positively associated with dEMG responses across all challenge conditions, including hypercapnia (Pearson, r=0.804, p=0.054), hypoxia (Pearson, r=0.459, p=0.36) and inspiratory restriction (Spearman correlation r=0.714, p=0.111). * p<0.05; ** p<0.01; *** p<0.001.

### Donor to Host Connectivity

Pseudorabies virus (PRV) was used to transynaptically trace donor-host connectivity at 1-M and 1-Y post-transplantation. Applying PRV to the diaphragm ipsilateral to injury/transplant, and allowing 3 days for transport, labels pre-motor spinal neurons associated with that network, including donor cells that have become integrated (Fortino et al., 2022). A difference in the number of PRV+ cells was readily apparent in representative images (Figure 5A, B) between 1-M and 1-Y. Quantification of PRV+ donor cells relative to the number of primary labeled cells (phrenic motor neurons) revealed significantly fewer PRV+ donor cells at 1-Y compared to 1-M (p=0.0002; Figure 5C), suggesting temporal decline in donor to host connectivity. Correlational analyses of donor-host connectivity and diaphragm activity revealed one statistically significant positive relationship between the number of PRV+ donor cells and diaphragm activity during hypercapnia at 1-Y time point (r=0.812, p=0.004; Figure 5E).

**Figure 5:**
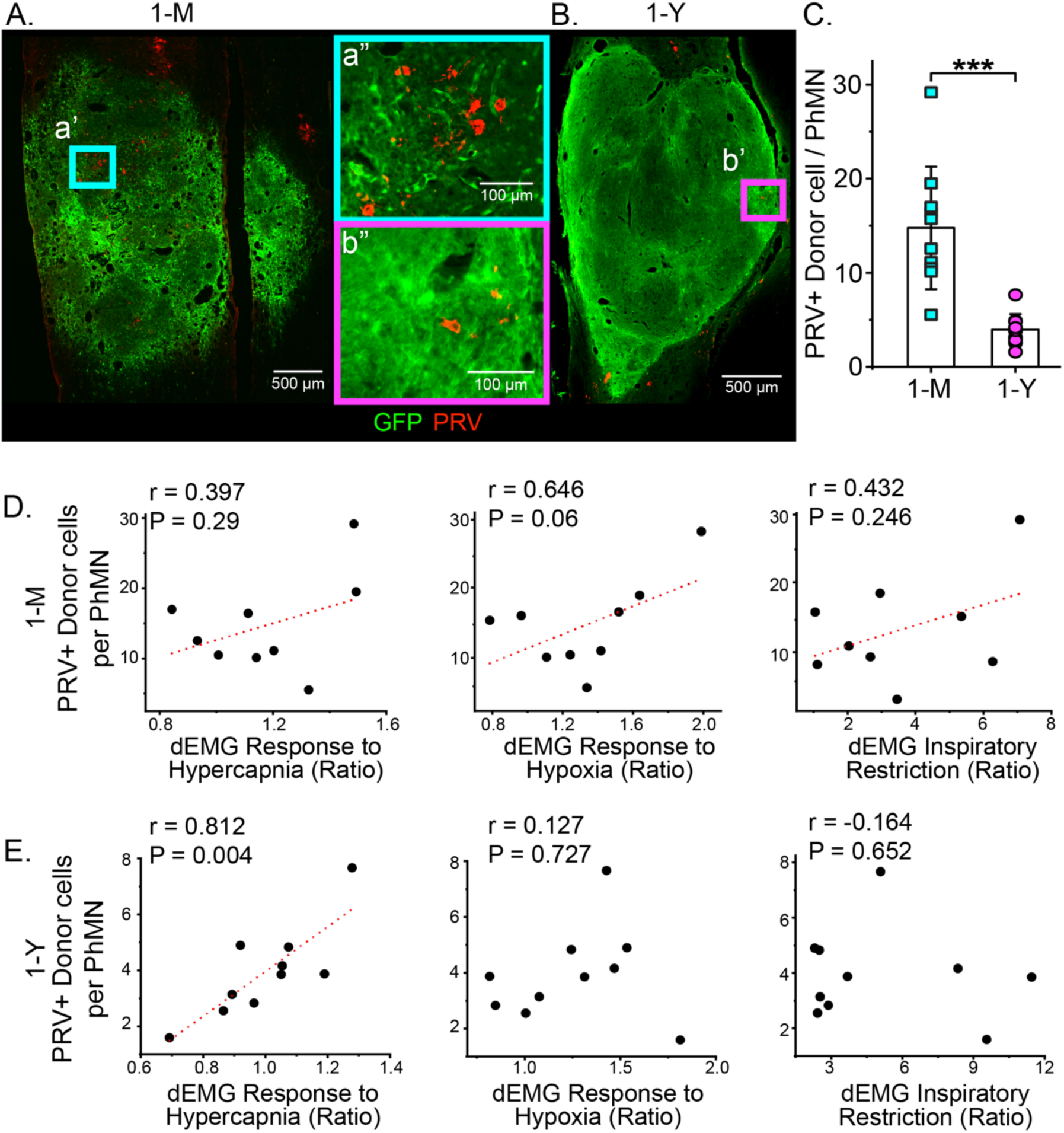
Fewer PRV-labeled donor neurons observed 1-Y post-transplantation. (A-B) Representative images of PRV-labeled donor-neurons at 1-month (1-M) and 1-year (1-Y) post-transplantation. Insets (a”, b”) show high-magnification views of PRV labeling within transplant. (C) Quantification of PRV+ donor neurons normalized to total phrenic motor neurons (PhMN) revealed less PRV-labeled neurons per motoneuron at 1-Y compared to 1-M (2-sample t-test, p=0.0002), suggesting a temporal decrease of donor-to-host connectivity. Correlational analyses at 1-M (D) and 1-Y (E) revealed a significant relationship between PRV-labeling and dEMG response to hypercapnia at 1-Y post-transplantation. * p<0.05; ** p<0.01; *** p<0.001.

## Host-Donor Serotonergic Innervation

Assessment of serotonergic fiber growth into transplanted tissue revealed varying degrees of serotonergic innervation to the transplant between animals. Qualitatively, the density of serotonin (5-HT) labeling in the more caudal regions of the transplant was greater in 1-Y animals (Figure 6A-B). Quantification of 5-HT-labeling revealed greater overall density (p=0.001; Figure 6C), and a greater percentage of 5-HT to GFP labeling in 1-Y transplant recipients compared to 1-M (p<0.0001; Figure 6D). Correlational analyses at 1-M (Figure 6E) and 1-Y (Figure 6F) revealed a significant relationship between relative 5-HT labeling and diaphragm EMG response to hypoxia at 1-Y post-transplantation.

**Figure 6.**
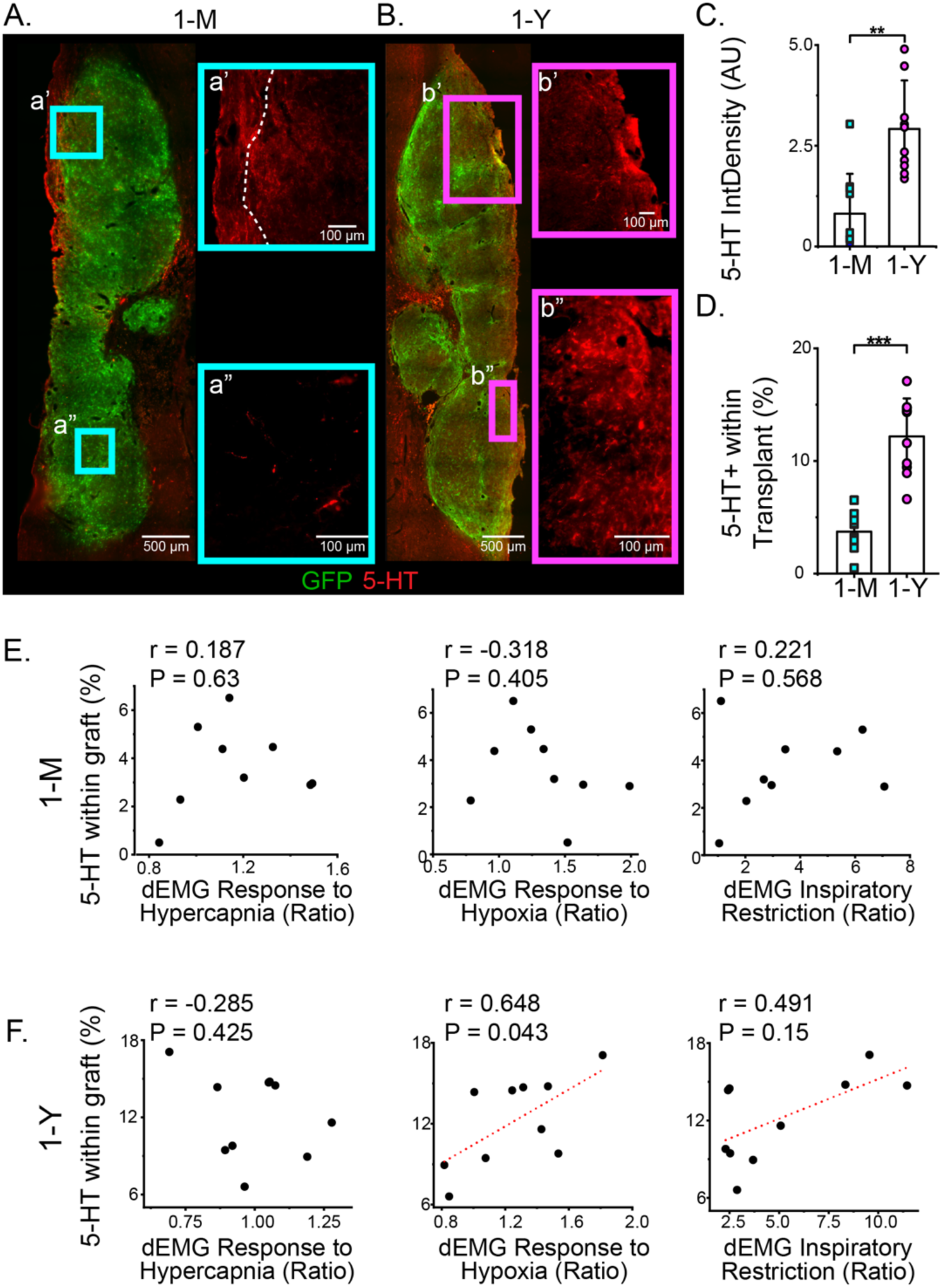
Serotonergic innervation of donor tissue is greater at 1-Y post-transplantation. (A, B) Representative images of serotonergic (5-HT) labeling within GFP⁺ transplants at 1 month (1-M) and 1 year (1-Y) post-transplantation. Insets (a′, b′) show comparable serotonergic fiber presence in rostral transplant regions, whereas caudal regions (a″, b″) demonstrate markedly higher 5-HT fiber density at 1-Y. (C) Quantification of 5-HT–associated integrated density was significantly greater in 1-Y transplants compared with 1-M (2-sample unpaired Student’s t test, p=0.0013). (D) The percentage 5-HT+ labeling within GFP+ transplant area (ROI) was significantly higher in 1-Y transplant recipients (unpaired Student’s t test, p<0.0001). Correlational analyses at 1-M (E) and 1-Y (F) revealed a significant relationship between 5-HT labeling and dEMG response to hypoxia at 1-Y post-transplantation. * p<0.05; ** p<0.01; *** p<0.001.

### Immune Cell Presence Increases Over Time

Immunohistochemistry for Iba1 was used to assess presence of microglia/macrophages within transplanted tissue at 1-M and 1-Y (Figure 7A, B). Iba1 immunoreactivity was quantified as mean pixel density within the GFP+ transplant-defined region of interest using ImageJ, consistent with intensity-based analyses used for serotonergic axons in prior studies (Starkey et al., 2012; Bartus et al., 2014). Qualitatively, Iba1+ cells exhibited heterogeneous morphologies throughout the transplant, including ramified, amoeboid, and rod-shaped profiles. Quantitative analysis revealed no significant difference in overall Iba1 immunoreactivity between 1-M and 1-Y transplant recipients, although greater inter-animal variability was observed at the 1-Y time point. Correlational analyses demonstrated a significant positive relationship between Iba1 immunoreactivity within the transplant and diaphragm EMG responses to inspiratory restriction at 1-Y post-transplantation (r=0.829, p=0.042; Figure 7F), whereas no such relationship was observed at 1-M (Figure 7E).

**Figure 7.**
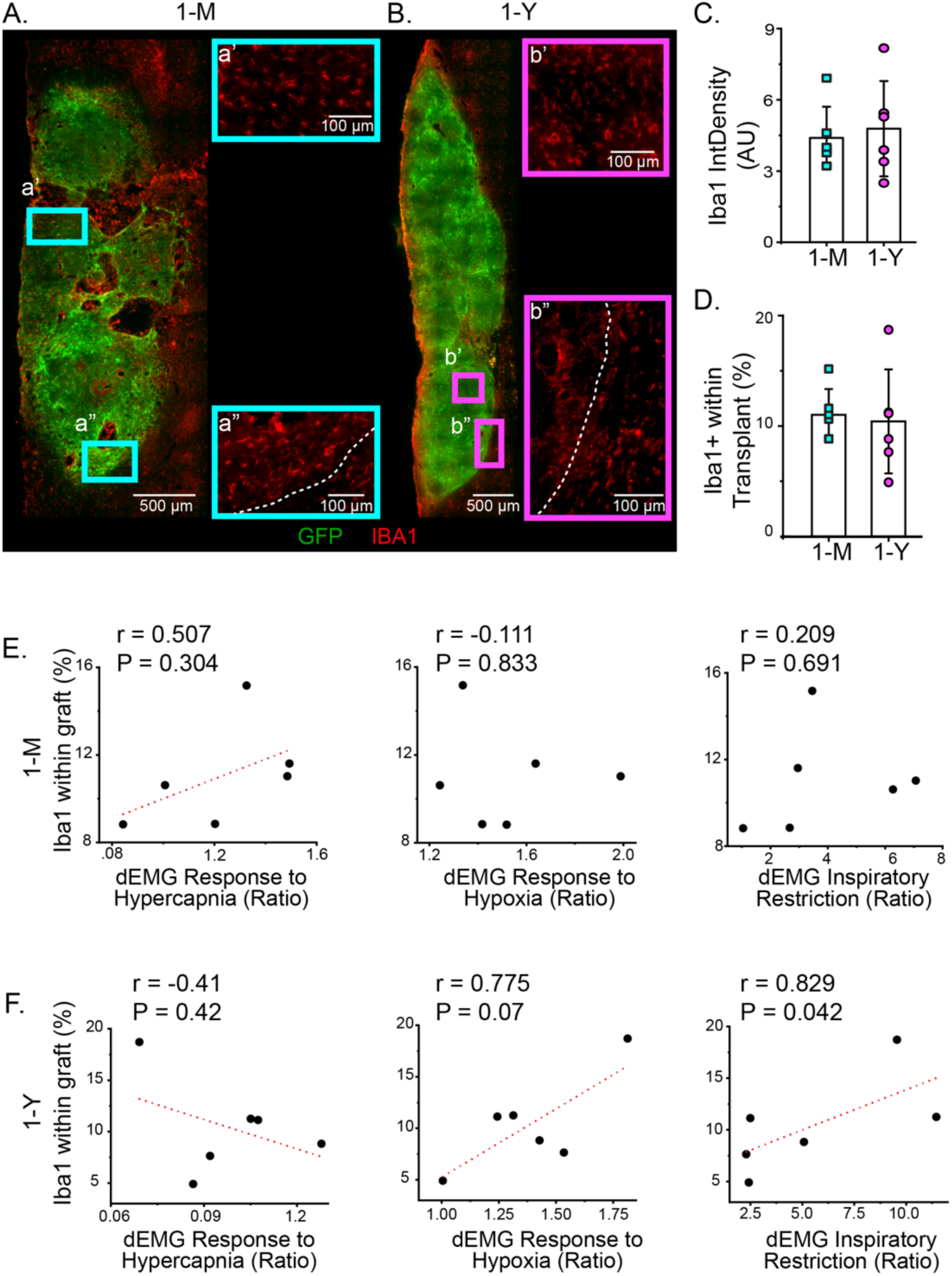
Microglial/macrophage (Iba1⁺) labeling persists over time. (A, B) Representative images of Iba1-labeling within GFP⁺ grafts at 1 month (1-M) and 1 year (1-Y) post-transplantation. Insets (a′, a″, b′, b″) show higher-magnification views of Iba1-labeling within distinct graft regions. (C, D) Quantification of Iba1+ integrated density (C) and percent of graft (GFP+) area that is positive for Iba1 labeling (D) revealed no significant differences between 1-M and 1-Y cohorts (unpaired Student’s t tests; p=0.7847 and p=0.7016, respectively). Correlational analyses at 1-M (E) and 1-Y (F) revealed a significant relationship between Iba1 labeling within transplant and dEMG response to inspiratory restriction at 1-Y post-transplantation.

## Discussion

This study provides a long-term evaluation of donor-host connectivity between transplanted cells and the spinal phrenic network following cervical SCI. By quantifying transneuronal labeling and diaphragm activity sub-acutely (1-month; 1-M) and chronically (1-year; 1-Y) after transplantation of embryonic spinal cord tissue, we identify a clear temporal shift in transplant-mediated repair: donor-host connectivity and respiratory recovery are robust in the subacute period but decline significantly at late chronic stages. Transneuronal tracing with PRV revealed significantly fewer labeled donor neurons at 1-Y, suggesting reduced connectivity between transplanted neurons and the phrenic motor network. Diaphragm activity in response to respiratory challenges was improved 1-M after transplantation, but was reduced at 1-Y with responses that no longer differed from injured control animals. These results reveal that embryonic spinal cord transplants possess the intrinsic capacity to reconstruct host circuitry and transiently improve respiratory activity. While transplanted cells survive chronically, long-term efficacy appears to be limited by progressive reduction of donor-host connectivity. These results highlight a critical consideration for successful long-term integration, emphasizing the need for additional strategies to guide, reinforce and/or retain donor-host connectivity.

These results build on decades of evidence that neural progenitor cells can integrate with host spinal networks and contribute to functional restoration (Reier, 1985; Bregman and Reier, 1986; Jakeman et al., 1989; Jakeman and Reier, 1991; Lu et al., 2003; Lepore et al., 2004; Lin et al., 2017). However, the majority of prior work has focused on acute and sub-acute times after transplantation, with limited effort to assess chronic outcomes. Notable examples of work that has assessed outcomes beyond a year post-transplantation are allogeneic developing cat spinal cord tissue (Reier et al., 1992) and xenogeneic human pluripotent stem cell-derived neural stem/progenitor cells (Lu et al., 2017; Zheng et al., 2023), that were transplanted into injured cat and rat spinal cord, respectively. While focused on locomotor network repair following thoracic SCI, both studies reported varying degrees of donor cell/tissue integration with the host. They also noted, however, that these donor tissues mature more slowly than allogeneic rat tissues used in the present work, so the time course of donor-host integration would likely not compare well to the present work and may require even longer-term studies to demonstrate progressive loss of connectivity. In contrast, Jakeman and Reier (1991) explored the integration of developing rat spinal cord into the injured lumbar spinal cord of immune competent rats (allografts), and found very limited donor-to-host integration beyond a year after transplantation in a small number of animals. Importantly, there did not appear to be continued growth to or from donor tissues, and if anything, the short distance integration seen >1 year (albeit in only a few animals) was less than seen at shorter time-points. Consistent with these previous studies, the present work also shows that the transplantation of neural progenitor cells from developing spinal cord tissue can contribute to repair of damaged neural networks, but consideration needs to be given to the extent of integration and whether it is retained long-term.

### Donor Tissue Composition

Transplants of developing spinal cord tissue comprise a variety of cell types including but not limited to maturing neurons (Diener and Bregman, 1998), glia (Miya et al., 1997), neural (neuronal and glial) progenitor cells (Spruance et al., 2018; Aceves et al., 2023), extracellular matrix (Wiese and Faissner, 2015), and resident immune cells (microglia) (Norris and Kipnis, 2019). Given that these donor tissues are developing, it is important to appreciate the range of cell phenotypes that survive transplantation, mature, and become integrated with the host over time. While one prior study found that intraspinal transplants into in-bred syngeneic mice did not show a significant temporal loss of neurons from 6 weeks to 6 months post-transplant (Baltazar et al., 2023), there was a reduction in the number of both NeuN and PRV positive donor cells in the present work from 1-M to 1-Y. This decline in the present work reflects the apparent loss of donor neurons (NeuN+) and donor-host connectivity (PRV+) one year following transplantation into the injured cervical spinal cord. Notably, the apparent loss of donor-host integration exceeded the apparent neuronal loss, suggesting a reduced synaptic connectivity among surviving cells. Despite the observed changes between 1-M and 1-Y, we also found that serotonergic innervation, Iba1 reactivity, and neuronal density correlated positively with functional outcomes in 1-Y transplant recipients. The contribution of each of these anatomical factors to functional outcomes over time should be explored further in future work.

Microglia and infiltrating macrophages play critical roles in both the initial and chronic phases of SCI (Ahuja et al., 2017; Bellver-Landete et al., 2019; Brockie et al., 2021; Brennan et al., 2022). These cells exist on a continuum from pro-inflammatory, synapse-pruning M1 phenotypes (Xu et al., 2021; Pereira-Iglesias et al., 2025) to neuroprotective M2 states (Guo et al., 2022). While chronic immune activation can persist for years after injury (Rezvan et al., 2020; Hellenbrand et al., 2021; Li et al., 2022), cellular therapies can modulate this response, promoting pro-reparative microglia that support neuronal homeostasis and recovery (Karova et al., 2019; Doulames et al., 2021; Delarue et al., 2025). Our results are consistent with this, revealing sustained Iba1+ labeling within the transplant at 1-Y. It is possible that such immune cell presence initially contributes to synaptic pruning or cytotoxicity (Zhang et al., 1997; Pereira-Iglesias et al., 2025), followed by a transition toward a more reparative M2-like state. Interestingly, elevated Iba1 density was positively correlated with improved responses to hypoxic and inspiratory restriction challenges, supporting a potential role for microglia in functional recovery through modulation of neuronal activity (Wang et al., 2022; Pereira-Iglesias et al., 2025). Future work could assess how combining anti-inflammatory strategies (e.g., pharmacologics, biomaterials, activity-based therapies) with transplantation could attenuate harmful inflammation and synergize with cell transplantation to influence long-term graft survival, connectivity, and repair.

Changes in NeuN and Iba1 labeling density must be interpreted cautiously, as density measurements made here reflect staining intensity within a defined region rather than absolute cell numbers. However, even without stereological cell counts or volumetric quantification, the observed differences likely reflect evolving tissue composition and microenvironmental changes within transplanted regions, consistent with the long maturational period of developing spinal tissue and the dynamic nature of chronic post-injury inflammation.

### Repairing the Phrenic Motor Network

The phrenic motor network represents one of the most clinically critical systems for evaluating functional neural repair, as even partial restoration of diaphragm control can markedly improve respiratory independence, quality of life, and survival after cervical SCI. Previous studies by our team (White et al., 2010; Lee et al., 2014; Spruance et al., 2018; Zholudeva et al., 2018; Zholudeva et al., 2024) and others (Lin et al., 2017; Sandhu et al., 2017), have also demonstrated that neural progenitor cell transplants can improve phrenic motor network repair and recovery after cervical SCI. Transplantation of neural progenitors into the lesion site 1-week following a mid-cervical contusion injury has been shown to increase the amplitude of diaphragm activity (measured by EMG) during baseline (breathing room air) conditions, 1-month after transplantation (Spruance et al., 2018; Zholudeva et al., 2018); Figure 2B-D). Although no significant differences were seen in the present work between injured and transplant-recipients at baseline and hypercapnic challenge, there were significant differences present under hypoxic and inspiratory restriction challenges. This suggests that donor cells contribute to phrenic network and diaphragm activity during this increased respiratory drive. Collectively, these studies show that integration of donor cells with the host phrenic network supports therapeutically mediated recovery. Thus, if the substrates contributing to the repair are disrupted over time there will be a concomitant decline in recovery.

To our knowledge, no studies to date have analyzed phrenic network repair or recovery of diaphragm activity 1-Y following neural cell transplantation. The present work has revealed that 1-Y post-transplantation, the degree of donor-host connectivity with the phrenic motor network, and the extent of recovered diaphragm activity under challenged conditions, was significantly less than seen sub-acutely after injury and transplantation. The improved ability of transplant recipients to respond to respiratory challenge (hypoxia and inspiratory restriction) that was seen 1-M after transplantation was no longer present at 1-Y. Notably, the decline in response to challenge across all groups – including uninjured age-matched control animals - could partially reflect age-related decline in respiratory function. However, given that the left-dorsal hemidiaphragm of naïve animals retained a stronger response to hypercapnic challenge than transplant-recipients at 1-Y suggests that there are aspects of recovery that were lost over this time that supersedes aging. While the present data comes from static data points, future studies could implant indwelling muscle electrodes to explore the temporal change in diaphragm activity pre-and post-SCI, and then weekly or monthly following transplantation.

An important consideration is that reduced PRV labeling does not necessarily equate to complete loss of connectivity. PRV uptake and transsynaptic transport depend on synaptic strength, activity, and physiological state. A reduced number of labeled donor neurons may therefore reflect weaker or less active synapses rather than their absence. If donor-host connections remain present but conduct signals less efficiently, the 72-hour tracing period used in the present work may be insufficient to detect them. Longer survival times or complementary tracers would help distinguish between loss of connections and reduced synaptic efficacy. Therefore, while the present results strongly suggest declining donor-host communication, they do not exclude the possibility that a subset of weakened, but still anatomically present connections persist into the chronic period.

Regardless, the extent of chronic donor-host connectivity reflected by PRV labeling, albeit limited, correlated strongly with diaphragmatic response to hypercapnic challenge. The stronger correlation seen at this later time point may reflect refinement of connectivity (e.g., synaptic pruning or strengthened connections) with networks involved in phrenic activity during exposure to hypercapnia. Given that the donor tissue comprises a vast range of spinal neurons, an important consideration is that not all cells necessarily benefit recovery (e.g. (White et al., 2010)). Accordingly, temporal changes in connectivity may select for cells that perform specific functions, or even limit the extent of phrenic recovery that can be achieved at this 1-year time point. This supports the notion that careful attention should be given to the donor phenotypes that are transplanted, survive, and integrate with networks that are being repaired (Zholudeva and Lane, 2019a, b, 2022b).

These results highlight the need for strategies that preserve or reinforce donor-host connectivity over extended periods. Transplanting dissociated embryonic tissue provides a rich substrate of neurons, progenitors, glia, and extracellular matrix capable of forming functional relays, but even these comprehensive donor tissues and newly formed donor-host networks appear vulnerable to loss over time. Transplanting cells without any other intervention relies on environmental cues from the injured host spinal cord to appropriately and consistently guide transplant integration. However, the injured spinal cord contains insufficient cues to promote appropriate integration, especially under chronic conditions. Combining other therapeutic strategies with cell transplantation may be needed to optimize lasting repair and recovery (Pieczonka and Fehlings, 2023; Jagrit et al., 2024). Approaches such as activity-based therapies, neuromodulatory stimulation, targeted delivery of trophic factors, or transplantation of defined neuronal subtypes may help strengthen and stabilize donor-derived relays. For instance, there is substantial evidence to show that activity-based therapies (e.g., intermittent hypoxia) can be used to entrain network plasticity and connections and enhance network function (Dougherty et al., 2018; Welch et al., 2020; Gonzalez-Rothi and Lee, 2021; Perim et al., 2021). Accordingly, pairing activity-based therapies with neural transplantation (Lu et al., 2022) may be one means of enhancing a greater and more consistent host-donor-host integration, and could be a viable means of retaining that connectivity over time.

### Closing Remarks

This paper represents a proof-of-principle for evaluation of transplant-host connectivity and its impact on recovery from SCI. Consistent with prior work, the results demonstrate that embryonic spinal cord transplants can restore meaningful network connectivity and improve phrenic function after cervical SCI. However, here we have shown that these improvements decline substantially by one-year post-injury and transplantation. This temporal decline in functional outcome coincides with altered donor-host connectivity and changes in tissue composition. The present results underscore both the potential and the constraints of transplant-mediated circuit repair and provide a critical foundation for developing strategies aimed at achieving long-lasting respiratory recovery.

## Acknowledgements

Funding sources include: National Institutes of Health grant T32 NS121768 (AH); National Institutes of Health grant F31 NS125975 (TF); National Institutes of Health grant R01 NS104291 (MAL); California Institute for Regenerative Medicine DISC2-14180 (LVZ); Lisa Dean Moseley Foundation (LVZ, MAL); National Institutes of Health Virus Center grant P40 OD010996. We thank John Houle, Tatiana Bezdudnaya, and Itzhak Fisher for technical assistance and advice, and Jennifer Dulin who provided comments on early drafts of the manuscript. PRV was produced and supplied by Dr David Bloom (University of Florida) and the Center for Neuroanatomy with Neurotropic Viruses (CNNV) as part of the NIH Virus Center grant (P40 OD010996). Primary antibodies to PRV were supplied by Princeton University as part of Virus Center funding (P40 RR018604), and in agreement with Drs. Lynn Enquist and Estaban Engel. Present address for Victoria Spraunce: Spruance and Associates, Jacksonville, FL, USA. Present address for Tara Fortino: University of British Columbia, and the International Collaboration on repair discoveries (ICORD), British Columbia, Canada

---

**Supplementary Figure 1.**
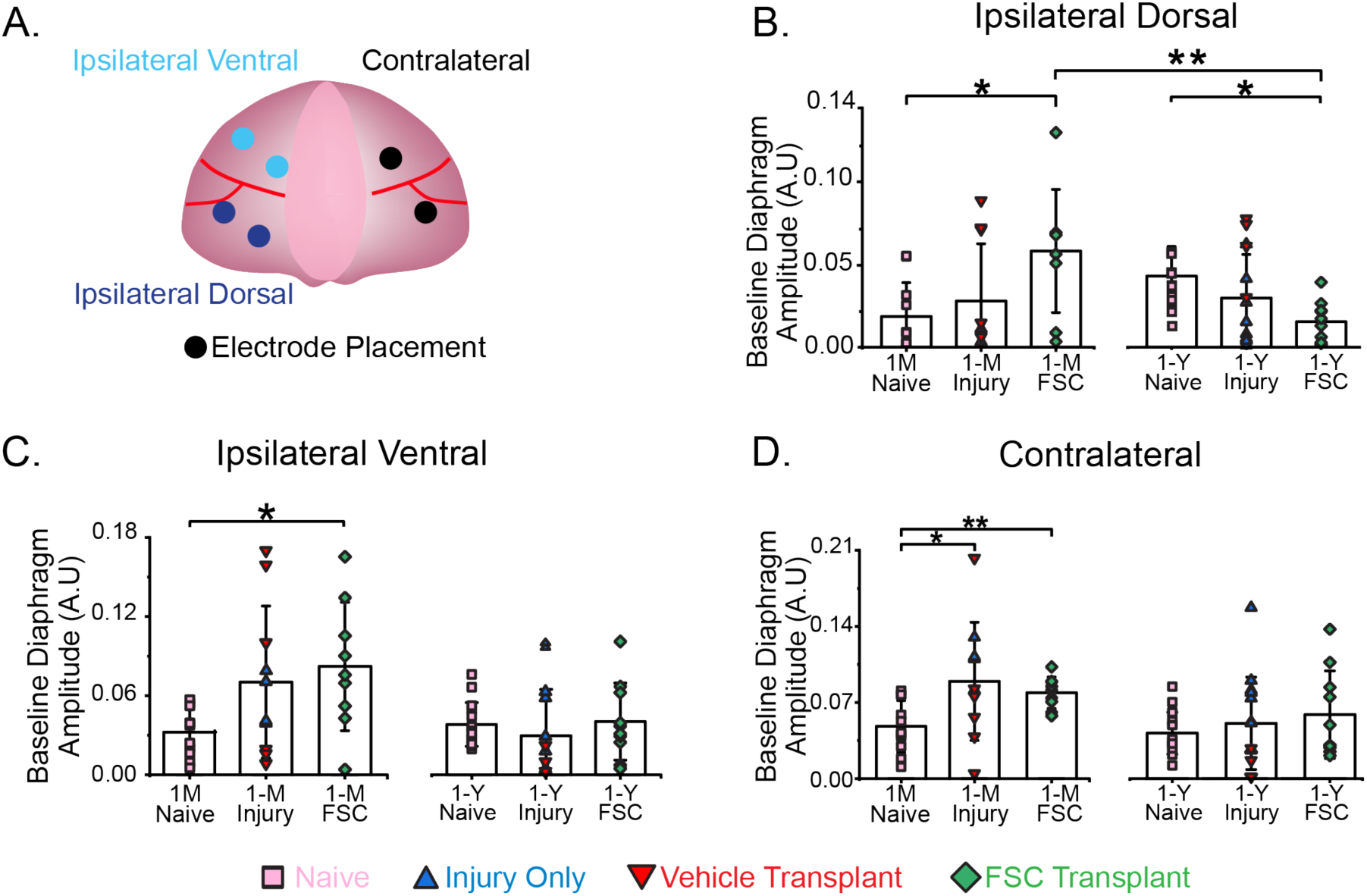
Diaphragm electromyography schematic and baseline amplitude across groups. (A) Schematic illustrating diaphragm electromyography (dEMG) recording regions: ipsilateral ventral (turquoise), ipsilateral dorsal (navy), and contralateral (orange). For ipsilateral dorsal and ventral recordings, electrode pairs were placed equidistant from each other within each region. Contralateral recordings were obtained using bipolar electrodes implanted dorsally and ventrally, separated by the midline blood vessel. (B) Baseline (eupneic) amplitude from the ipsilateral dorsal diaphragm in 1-month (1-M) and 1-year (1-Y) cohorts. Transplant recipients showed higher baseline activity compared with naïve animals at 1-M and 1-Y (p=0.003; p=0.030, respectively). Comparing transplant recipients at 1-M and 1-Y revealed a significant difference in ipsilateral dorsal dEMG (p=0.003). (C) 1-M transplant recipients showed significantly elevated baseline ipsilateral ventral dEMG activity compared with naïve controls (p=0.046). (D) 1-M injury and transplant recipients showed significantly elevated contralateral dEMG activity relative to naïve controls (p=0.032). Across panels, analyses were conducted using one-way ANOVA or Kruskal–Wallis ANOVA with appropriate post hoc testing, with significance defined as p<0.05. * p<0.05; ** p<0.01; *** p<0.001.

**Supplementary Figure 2:**
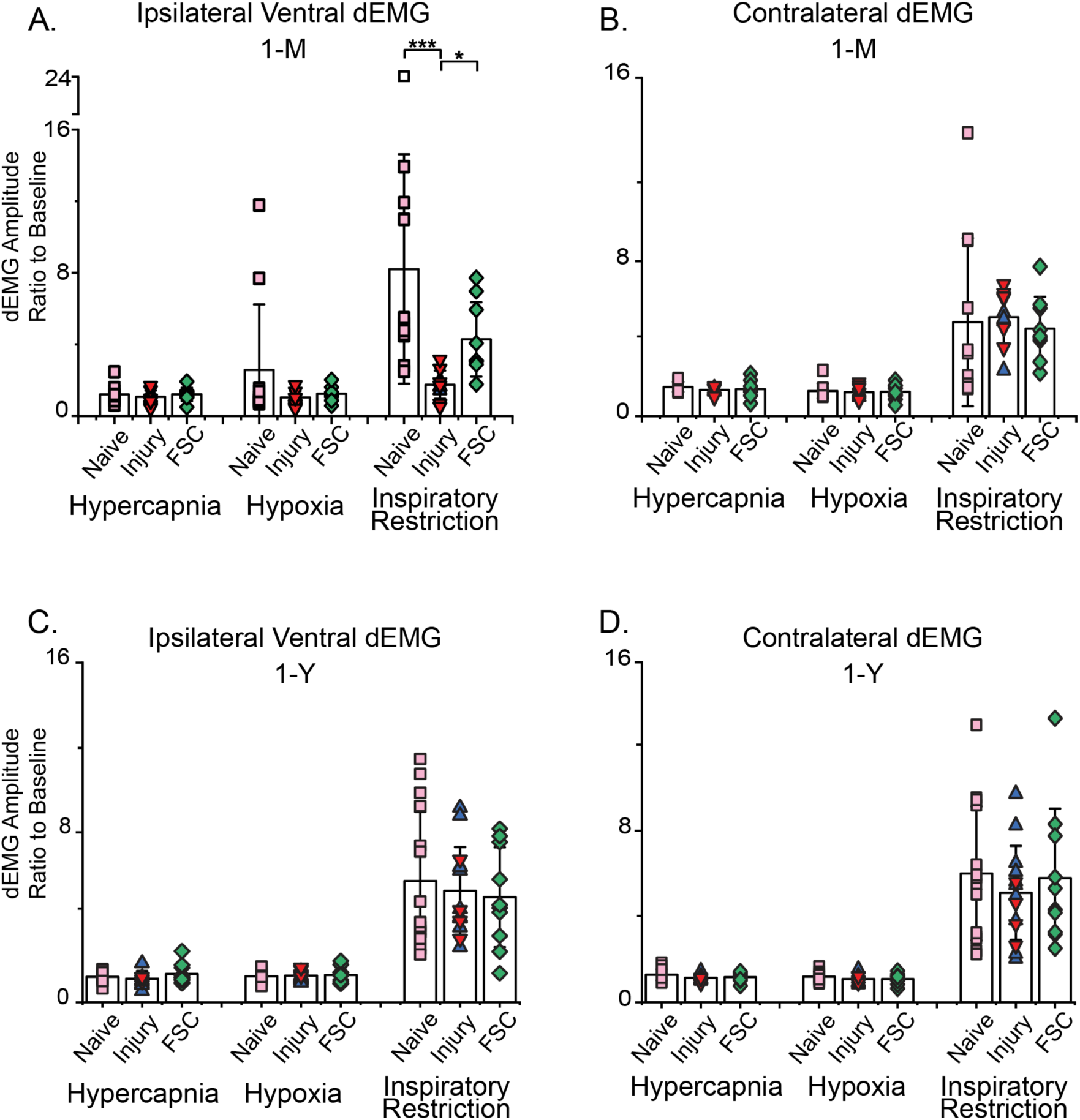
Response in diaphragm activity to respiratory challenges. (A) Ipsilateral ventral diaphragm activity during inspiratory restriction (as ratio to baseline) was significantly smaller in 1-M injured controls compared to naïve animals (p<0.0001). 1-M transplant recipients showed significantly higher dEMG compared to injured untreated animals (p=0.037). No other significant differences were observed at either 1-M (B) or 1-Y (C-D). Across panels, analyses were conducted using two-way ANOVA with appropriate post hoc testing, with significance defined as p<0.05. * p<0.05; ** p<0.01; *** p<0.001.

**Supplementary Figure 3.**
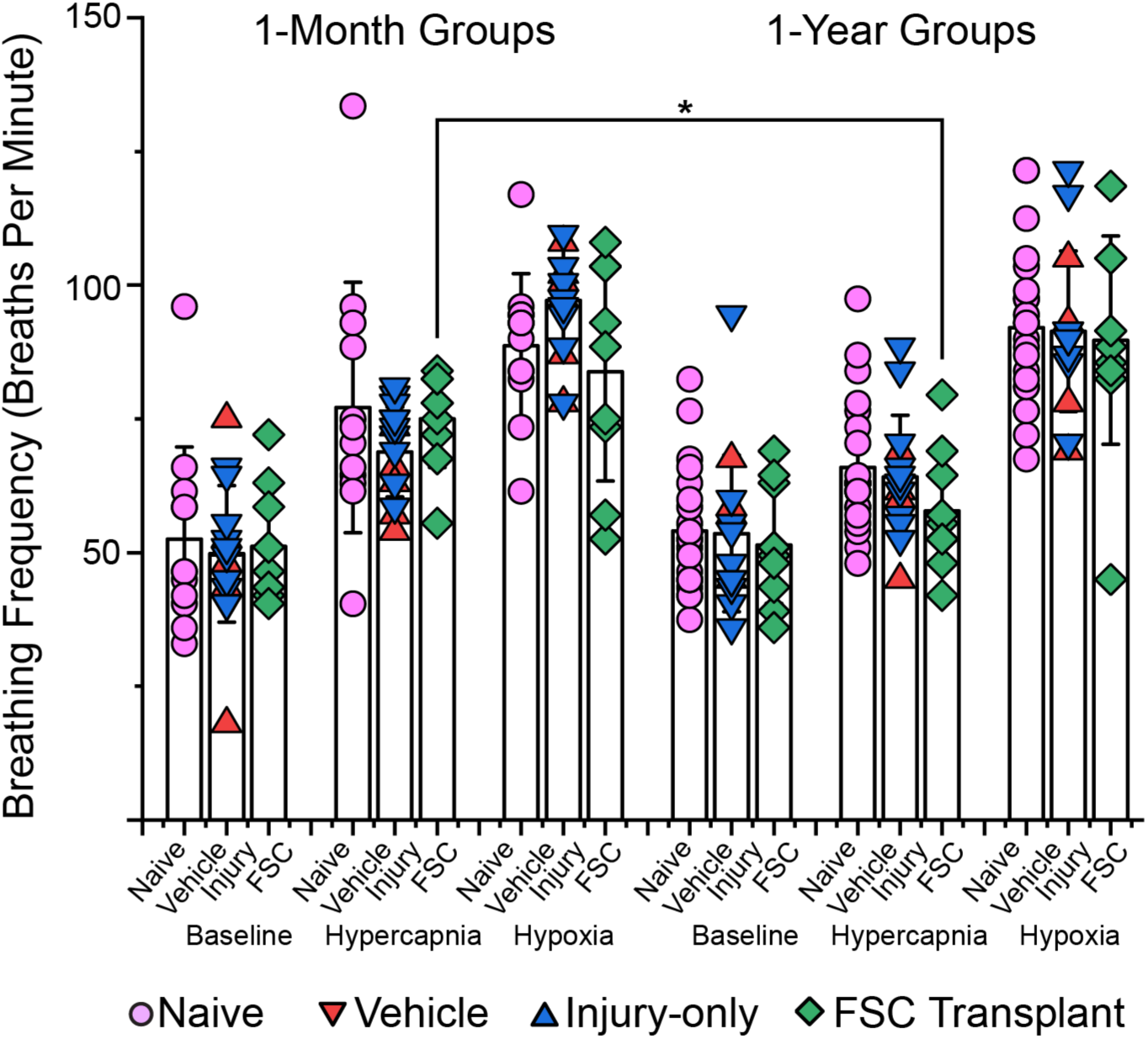
Respiratory frequency responses to chemoreflex challenges at 1 month and 1-year post-transplantation. Breathing frequency (breaths per minute) was measured during terminal dEMGs under baseline, hypercapnic, and hypoxic conditions at 1 month and 1-year post-transplantation. Animals were assigned to four groups: Naïve, vehicle (includes injury only), and FSC transplant. Individual data points represent single animals; bars indicate group means ± SD. Bracket indicates a significant group effect (p<0.05) between breathing frequency of FSC-recipients at 1-month compared to 1 year group. Statistical comparisons were performed using two-way mixed-measures ANOVA with Tukey post hoc testing, with significance defined as p < 0.05. * p<0.05; ** p<0.01; *** p<0.001.

**Supplementary Figure 4.**
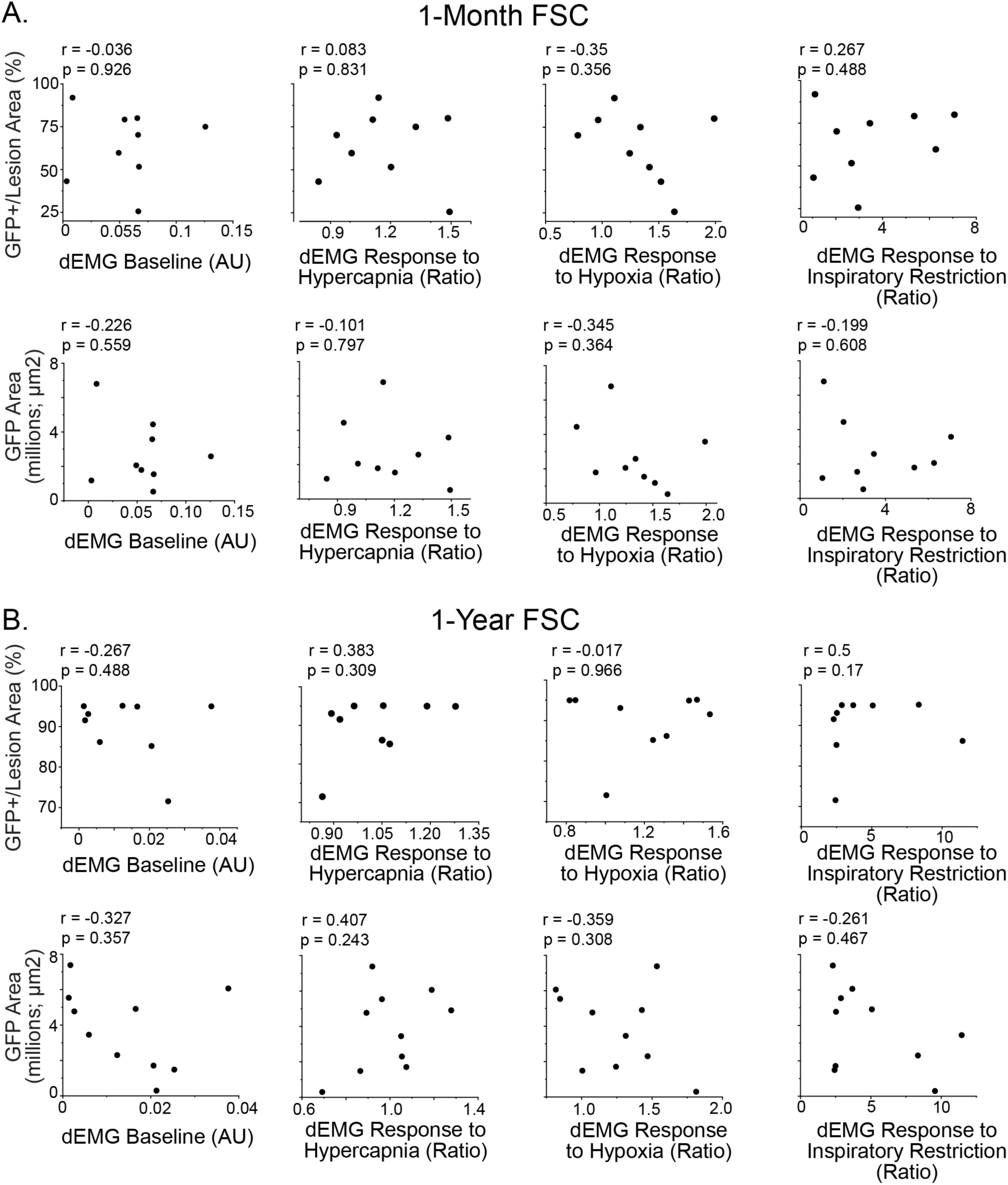
Donor graft size does not correlate with diaphragm EMG outcomes in FSC transplant recipients. (A) Correlational analyses between donor graft size and diaphragm electromyography (dEMG) in 1-month FSC transplant recipients. Top row shows correlations between GFP+ area normalized to lesion area (%) and dEMG baseline activity, hypercapnic response, hypoxic response, and inspiratory restriction response. Bottom row shows correlations between absolute GFP+ area (millions of µm²) and the same dEMG measures. No significant correlations were observed between graft size and any dEMG outcome at 1-month post-transplantation. (B) Correlational analyses between donor graft size and dEMG measures 1-year post-FSC transplantation, displayed as in (A). Neither normalized nor absolute GFP+ graft area significantly correlated with baseline diaphragm activity or responses to hypercapnic, hypoxic, or inspiratory restriction challenges at 1-year post-transplantation. Each data point represents a single animal. Pearson correlation coefficients (r) and corresponding p values are indicated within each plot. Statistical significance was defined as p<0.05.

